# Ultrasensitive capture of human herpes simplex virus genomes directly from clinical samples reveals extraordinarily limited evolution in cell culture

**DOI:** 10.1101/330860

**Authors:** Alexander L. Greninger, Pavitra Roychoudhury, Hong Xie, Amanda Casto, Anne Cent, Gregory Pepper, David M. Koelle, Meei-Li Huang, Anna Wald, Christine Johnston, Keith R. Jerome

## Abstract

Herpes simplex viruses (HSV) are difficult to sequence due to their large DNA genome, high GC content, and the presence of repeats. To date, most HSV genomes have been recovered from culture isolates, raising concern that these genomes may not accurately represent circulating clinical strains. We report the development and validation of a DNA oligonucleotide hybridization panel to recover near complete HSV genomes at abundances up to 50,000-fold lower than previously reported. Using copy number information on herpesvirus and host DNA background via quantitative PCR, we developed a protocol for pooling for cost-effective recovery of more than 50 HSV-1 or HSV-2 genomes per MiSeq run. We demonstrate the ability to recover >99% of the HSV genome at >100X coverage in 72 hours at viral loads that allow whole genome recovery from latently-infected ganglia. We also report a new computational pipeline for rapid HSV genome assembly and annotation. Using the above tools and a series of 17 HSV-1-positive clinical swabs sent to our laboratory for viral isolation, we show limited evolution of HSV-1 during viral isolation in human fibroblast cells compared to the original clinical samples. Our data indicate that previous studies using low passage clinical isolates of herpes simplex viruses are reflective of the viral sequences present in the lesion and thus can be used in phylogenetic analyses. We also detect superinfection within a single sample with unrelated HSV-1 strains recovered from separate oral lesions in an immunosuppressed patient during a 2.5-week period, illustrating the power of direct-from-specimen sequencing of HSV.

**Importance:** Herpes simplex viruses affect more than 4 billion people across the globe, constituting a large burden of disease. Understanding global diversity of herpes simplex viruses is important for diagnostics and therapeutics as well as cure research and tracking transmission among humans. To date, most HSV genomics has been performed on culture isolates and DNA swabs with high quantities of virus. We describe the development of wet-lab and computational tools that enable the accurate sequencing of near-complete genomes of HSV-1 and HSV-2 directly from clinical specimens at abundances >50,000-fold lower than previously sequenced and at significantly reduced cost. We use these tools to profile circulating HSV-1 strains in the community and illustrate limited changes to the viral genome during the viral isolation process. These techniques enable cost-effective, rapid sequencing of HSV-1 and HSV-2 genomes that will help enable improved detection, surveillance, and control of this human pathogen.

## Introduction

Herpes simplex virus-1 (HSV-1) and herpes simplex virus-2 (HSV-2) are alphaherpesviruses causing over 4 billion human infections that can manifest as oral and genital ulcerations, neonatal disease, herpetic keratitis, and encephalitis (1, 2). While HSV-2 has traditionally been associated with genital herpes, HSV-1 comprises the majority of first episode genital herpes infections in high income countries (3). HSV genome evolution is notable for extensive HSV-1 recombination within HSV-2 genomes, with no detectable HSV-2 recombination into HSV-1 genomes (4, 5).

To date, most human herpes simplex virus genome sequencing has been performed on culture isolates (6–10). Culture is a pragmatic method to enrich for viral sequences and many clinical virology labs have rich banks of cultured HSV isolates. However, without the ability to compare these sequences to sequence recovered directly from clinical samples, interpretation of sequence results has been tempered by the concern that culture isolates might not accurately represent viral sequence in vivo. Other viruses such as influenza and parainfluenza viruses have shown that culture adaptation results in radically different viral sequence and receptor binding properties that do not accurately reflect selection pressures in vivo (11–13). Culture of the polyomaviruses BK and JC viruses is often performed in SV40 large T-antigen immortalized cell lines, allowing near complete loss of the BKV and JCV large T-antigen via transcomplementation, representing loss of one-third of the viral genome (14, 15). Culture adaptation of human herpesvirus 6A/B results in large tandem repeats in the origin of replication and other regions that are not found in low-passage clinical isolates and likely help accelerate viral replication in vitro (16–18). Similarly, laboratory passage of human cytomegalovirus, Epstein-Barr virus, and varicella-zoster virus can result in surprisingly large deletions comprising multiple genes and kilobases (19–22).

Many clinical studies of HSV conducted at our institution and throughout the world have utilized swabs to obtain DNA, which have the advantages of being easily collected, stable at room temperature, and can be sequenced directly from the patient. To fully take advantage of the rapidly growing field of genomics to understand HSV pathogenesis and diversity, we created a high-throughput method for sequencing HSV from DNA swab and culture material. Capture sequencing has become commonly used in human exome sequencing, oncology panels, and for other herpes viruses (23–25). We report here the development of wet-lab and dry-lab tools for sequencing of HSV-1 and HSV-2 genomes directly from clinical specimens using a custom oligonucleotide hybridization panel. In our hands, these methods extended the range of HSV-1 and HSV-2 viral abundances from which whole genome recovery is possible by up to 5 logarithms. By recovering HSV-1 sequence direct from clinical specimens, we compare sequences from HSV-1 in clinical samples with clinical isolates recovered from culture on human fibroblast cells. We show extraordinary limited evolution of HSV-1 genomes during viral isolation. As an example of the power of our approach, we also report the first genomic detection of HSV-1 superinfection from a single oral swab.

## Results

### Development of standard operating procedure for HSV genome capture

To recover whole genomes directly from clinical swabs, we designed a specialized capture sequencing workflow for clinical HSV genomics. DNA is extracted from clinical swabs collected in universal transport media or proteinase K buffer and total DNA is quantitated (Figure 1A). HSV and beta-globin copy number are quantitated using specific qPCR.

**Figure 1.**
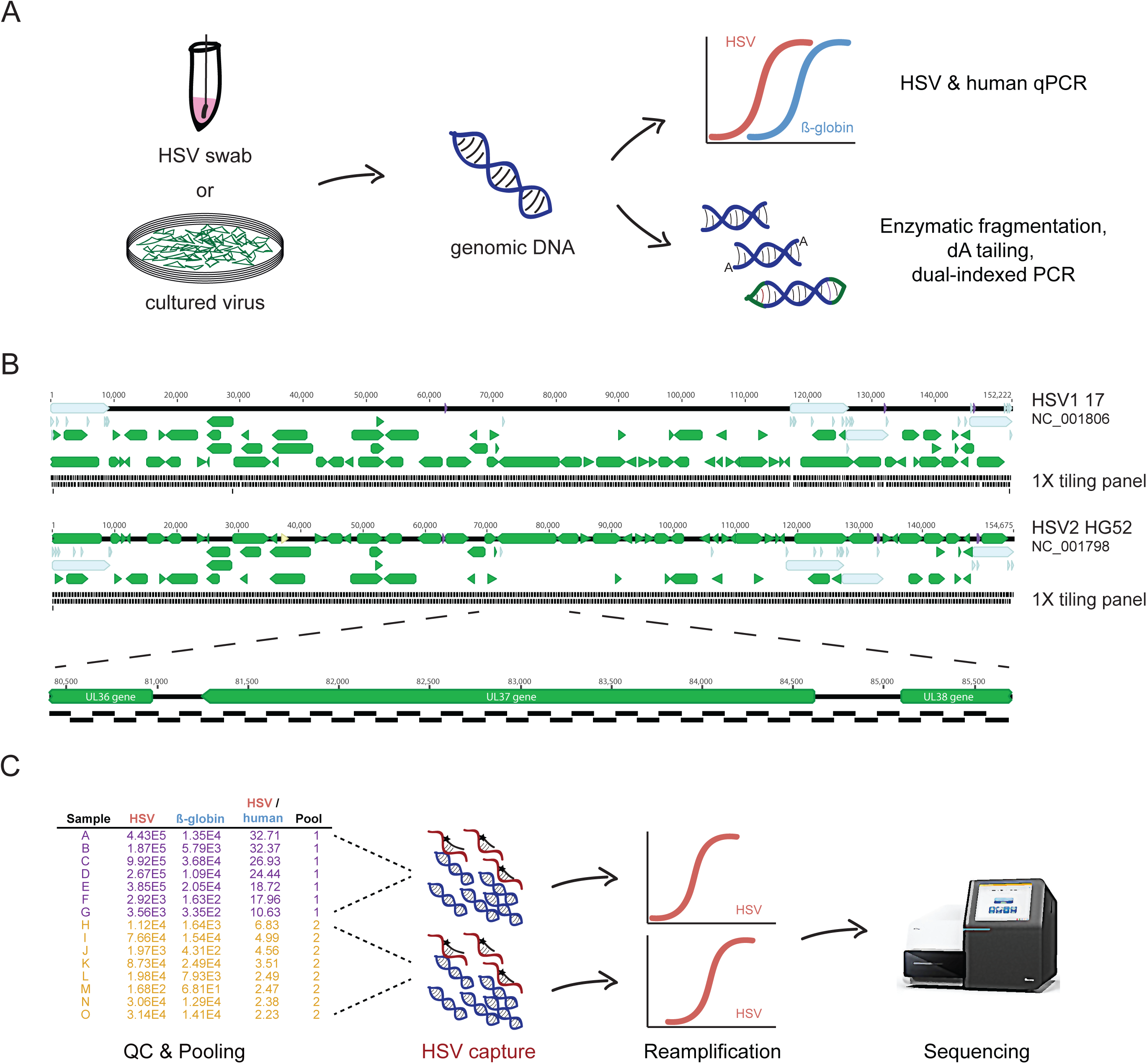
Experimental protocol. A) DNA is extracted from either clinical swabs in proteinase K buffer or cell culture supernatant. DNA is quantitated for HSV and beta-globin and it is enzymatically fragmented, end-repaired, dA-tailed, and TruSeq Y-adapters are ligated on. B) Design of 1X 120-bp tiling panel across HSV-1 and HSV-2 genomes. C) Samples are pooled in sets of 4-10 based on the HSV/beta-globin ratio to minimize variance in viral concentration and readjusted based on the total HSV copies present in each sample. D) A typical high concentration HSV-2 sample is shown (10^6^ copies/mL virus) based on shotgun sequencing and capture sequencing, illustrating a >3000-fold increase in viral sequence enrichment.

Based on our experience with the limited sensitivity of shotgun sequencing directly from HSV-2 clinical swabs, we developed a custom tiling oligonucleotide panel for HSV-2 based on the HG52 reference genome (NC_001798) (Figure 1B) (9). Experiments showed that while the HSV-2 capture panel could readily recover near complete genomes from HSV-2 material, it could only recover less than 30% of the HSV-1 genome from HSV-1 culture specimens (Figure S1A). Recovered regions of HSV-1 correlated with its average pairwise identity to HSV-2, requiring >85% pairwise identity for high coverage (Figure S1B). We thus designed an additional HSV-1 capture panel for subsequent HSV-1 capture experiments (Figure 1B).

To increase the cost-effectiveness of capture sequencing, we developed a pooling scheme for performing capture on dual-indexed libraries. While pooling schemes are common in many capture sequencing protocols, dealing with potential billion-fold differences in copy number between different HSV specimens along with differences in host background and variance of quantitation by qPCR required a different approach (Figure 1C). For example, inclusion of a high HSV copy number specimen in the same pool with a low copy number specimen could result in few reads being obtained for the lower copy number specimen, thus requiring re-enrichment of the low copy number library. Our protocol ranks libraries by the relative amounts of HSV and beta-globin present and pools of 5-10 specimens are chosen based on the variance of HSV/human ratio present in the samples prepared. We generally prepare 30-50 pre-capture libraries in batch, resulting in approximately 4-7 pools for capture. Samples in pools may be subsequently reassigned to a different pool based on the total copy number of HSV-1 and HSV-2 present. Because different amounts of HSV are present in each pool, we perform the post-enrichment amplification step with an initial 10 cycles of PCR followed by monitoring of additional cycles on lower concentration pools by either SYBR-Green or iterative checking by agarose gel electrophoresis to a maximum of 20 PCR cycles after capture. Pools are sequenced on 2×300 bp Illumina MiSeq runs to enhance recovery of particularly high GC regions and the multiple repeats present in HSV genomes. The finished protocol as illustrated on culture specimens results in significant enrichment across the HSV-2 genome (Figure 1D).

### Development of a custom pipeline for HSV assembly and annotation

We developed a computational pipeline (Figure 2) to rapidly extract and annotate near-full length HSV genomes from raw Illumina sequencing reads. By employing a combination of reference-guided and assembly-based methods to construct consensus sequences, we were able to recover up to 99% of the genome.

**Figure 2.**
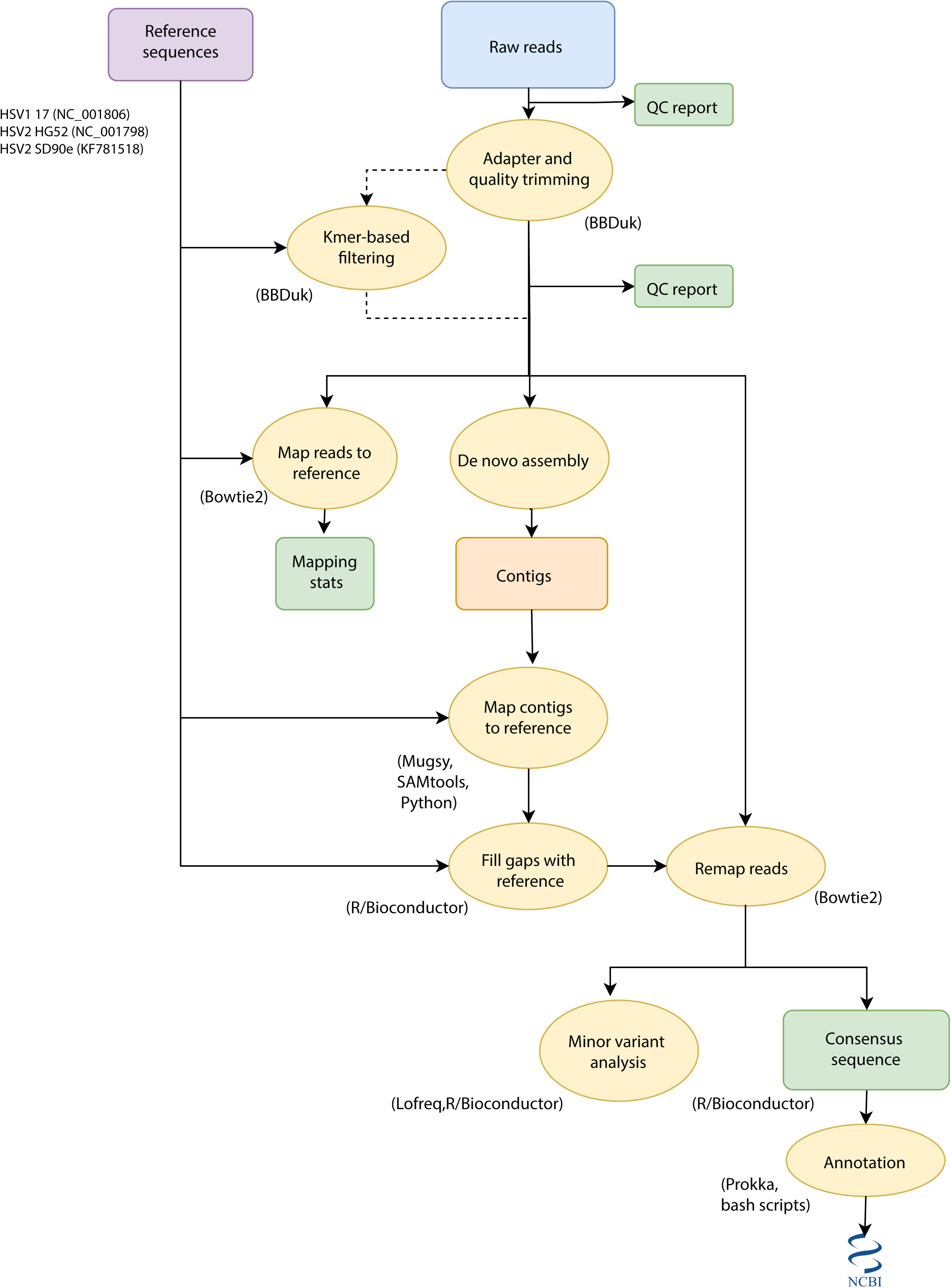
Overview of pipeline for assembly and annotation of HSV sequences. Raw reads are adapter and quality trimmed using BBDuk. If pre-capture shotgun HSV libraries are sequenced, trimmed reads are subjected to k-mer filtering prior to assembly to prevent tedious assembly of the human genome. Reads are de novo assembled using SPAdes v3.11 and mapped to each of three reference genomes to determine whether HSV-1 or HSV-2 was sequenced. Contigs are mapped to the chosen reference and gaps are filled with reference sequence. Finally, reads are mapped to this sequence in order to determine the consensus sequence before annotation and submission to NCBI.

The workflow starts with quality analysis of raw reads followed by trimming to remove adapters and low-quality regions. For samples sequenced without target capture enrichment or with a low percentage of HSV reads, a k-mer based filtering method is used to enrich for HSV reads based on similarity to the HSV-1 strain 17 and HSV-2 strain HG52 and SD90e reference sequences (Figure 2). The removal of off-target reads significantly speeds up downstream processing steps by preventing *de novo* assembly of mammalian genomes. Preprocessed reads are *de novo* assembled into contigs and the reference sequence is used to order these contigs and fill in any gaps. Reads are then mapped to this resulting template and custom scripts are used to construct the final consensus sequence. Finally, the consensus sequence is annotated and prepared for Genbank deposit. Our pipeline combines several previously published open-source tools with custom scripts and can be run on desktop computers, servers and high-throughput computing clusters. On average, a single sample containing about 700,000 raw reads run on a machine with 14 cores takes about 15 minutes.

### Accuracy of capture-based sequencing

To validate the accuracy of our sequencing method, we compared thymidine kinase (UL23) sequences obtained from PCR-Sanger sequencing and those obtained from our capture sequencing method for eight strains of HSV-1 and eight strains of HSV-2 (Figure S2). For Sanger sequencing, UL23 was PCR amplified from genomic DNA and Sanger sequenced to a minimum of 2X coverage. For the WGS genes, majority consensus sequence for the UL23 CDS was extracted from the annotated assembly and aligned against the corresponding Sanger sequence. No consensus variants were recovered from either of the two genes in either HSV-1 or HSV-2, yielding an accuracy of 100%.

### Limits of genome recovery

To determine the lower limit of capture for our whole genome sequencing method and to understand the determinants of our on-target percentage and coverage statistics, we performed capture sequencing on HSV-1 and HSV-2 clinical samples across a range of concentrations (Figure 3, Table S1). We calculated the pre-capture ratio of HSV mass to human DNA mass based on the quantities of HSV and beta-globin recovered in the initial qPCR reaction. We then compared the pre-capture HSV mass ratio to the on-target fraction of HSV reads after the capture as a proxy for genome recovery.

**Figure 3.**
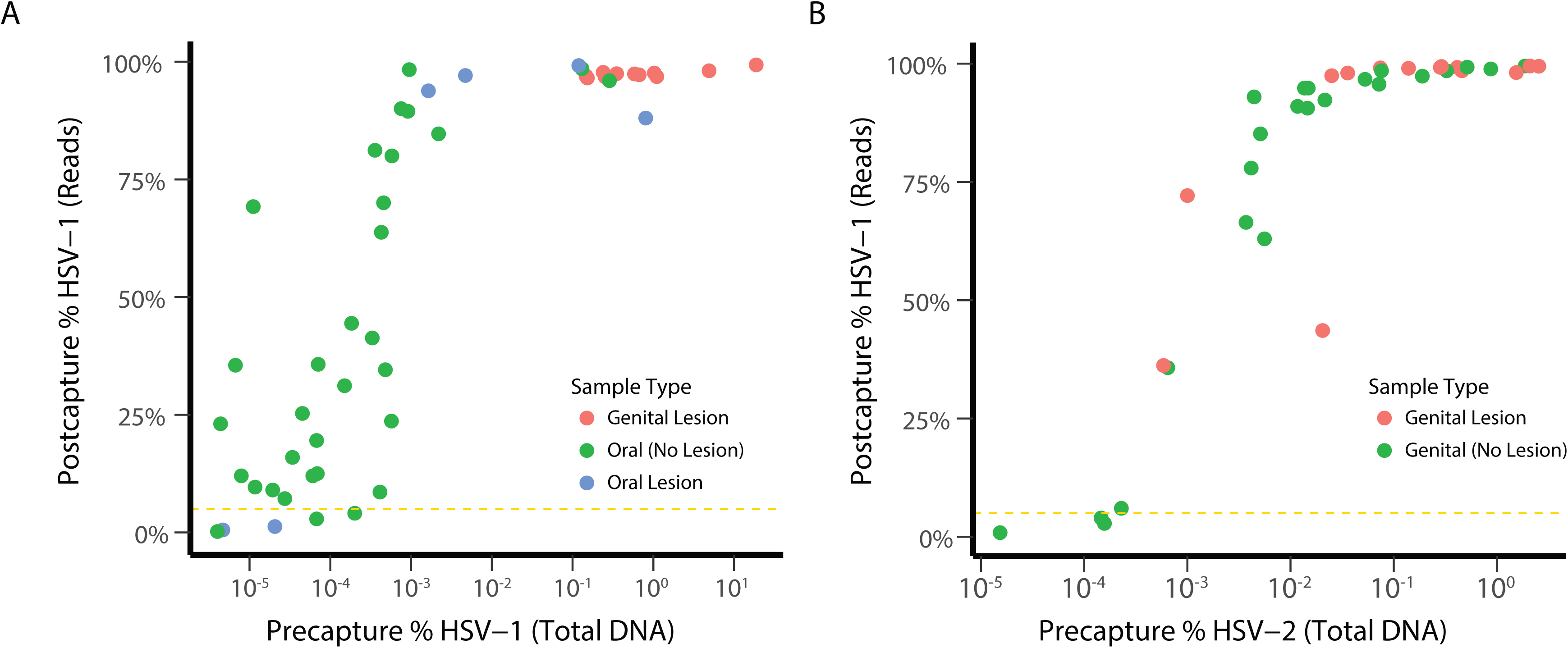
Capture sequencing allows near complete genomes from all symptomatic HSV clinical samples. Efficiency of sequence enrichment from clinical samples for A) HSV-1 and B) HSV-2 is depicted. Precapture HSV percentage of total DNA is shown on the X-axis based on qPCR values for HSV and beta-globin. Postcapture HSV percentage is shown on the Y-axis based on percent of total reads mapping to HSV (on-target percentage). Sample types are labeled by color for genital lesion (red), oral lesion (blue), asymptomatic oral shedding (green), or asymptomatic genital shedding (light green). The gold dotted line denotes 2% post-capture HSV reads, above which near complete genomes were obtained.

We find on average 10,000X enrichment of viral sequences with our capture panels with a maximum of 100,000X (Figure 3). With this approach we have recovered whole genomes from HSV-1/2 samples with viral loads lower than 10^2^ copies/rxn. Using an arbitrary cut-off of 5% on-target fraction of post-capture HSV reads, we can recover genomes from pre-capture ratios of 10^−7.40^ for HSV-1 and 10^−5.78^ for HSV-2, corresponding to approximately 10^3^ copies/mL for HSV-1 and 10^4^ copies/mL for HSV-2. Based on these copy numbers, we calculate that with capture sequencing we will be able to recover whole HSV genomes directly from nearly all swabs obtained by our clinical lab for symptomatic lesions and approximately 85% of HSV-positive swabs from asymptomatic persons for clinical studies (26).

### Sequencing of culture versus clinical specimens in HSV1

With the ability to recover whole HSV genomes directly from clinical specimens, we sought to address to what extent does sequence obtained from HSV-1 isolates obtained during routine culture in our clinical virology lab reflect viral sequence present in clinical swabs? We obtained 17 pairs of original clinical HSV-1 swabs in universal transport media that had associated positive HSV-1 culture results on human fibroblast cells. These HSV-1 isolates were derived from a variety of specimens, including bronchoalveolar lavage, oral swabs, vaginal swabs, and penile swabs (Table S2). All HSV-1 isolates were in culture for fewer than 7 days (range of 2-7 days) and only one isolate (sample G9) was passaged after isolation.

We sequenced these samples to a median of 547,494 reads (IQR 352,038-830,777; n = 34) and we recovered near-full length consensus genomes from as low as 101,000 reads. Median coverage was 518x (IQR 276-741x; n = 34) with up to 99.6% of quality and adapter-trimmed reads being on-target for HSV-1 (median 99.2%, IQR 99.0-99.3, n = 34).

HSV-1 UL and US sequences recovered directly from clinical specimens were nearly identical to those recovered after isolation from human fibroblasts (Figure 4). Allowing for all mutations, UL-US culture pairs had on average 20 SNVs (range 2-59), and most of these were present in repetitive elements in genes US12 and US5 that likely represent sequencing/assembly artifacts. After accounting for missing data (N’s), homopolymers (>8 nucleotides), and sequencing/assembly artifacts due to difficult loci such as high GC repeats in UL36, US5, and US12 genes, 14 of the 17 pairs of specimens were entirely identical in the UL-US region. One verified mutation was recovered in sample pair H5, with a synonymous C->T mutation in the consensus sequence at nucleotide 603 in UL39. The original H5 sample had a 55% C, 45% T allele frequency at the locus, while the culture sample was 4% C, 96% T.

**Figure 4.**
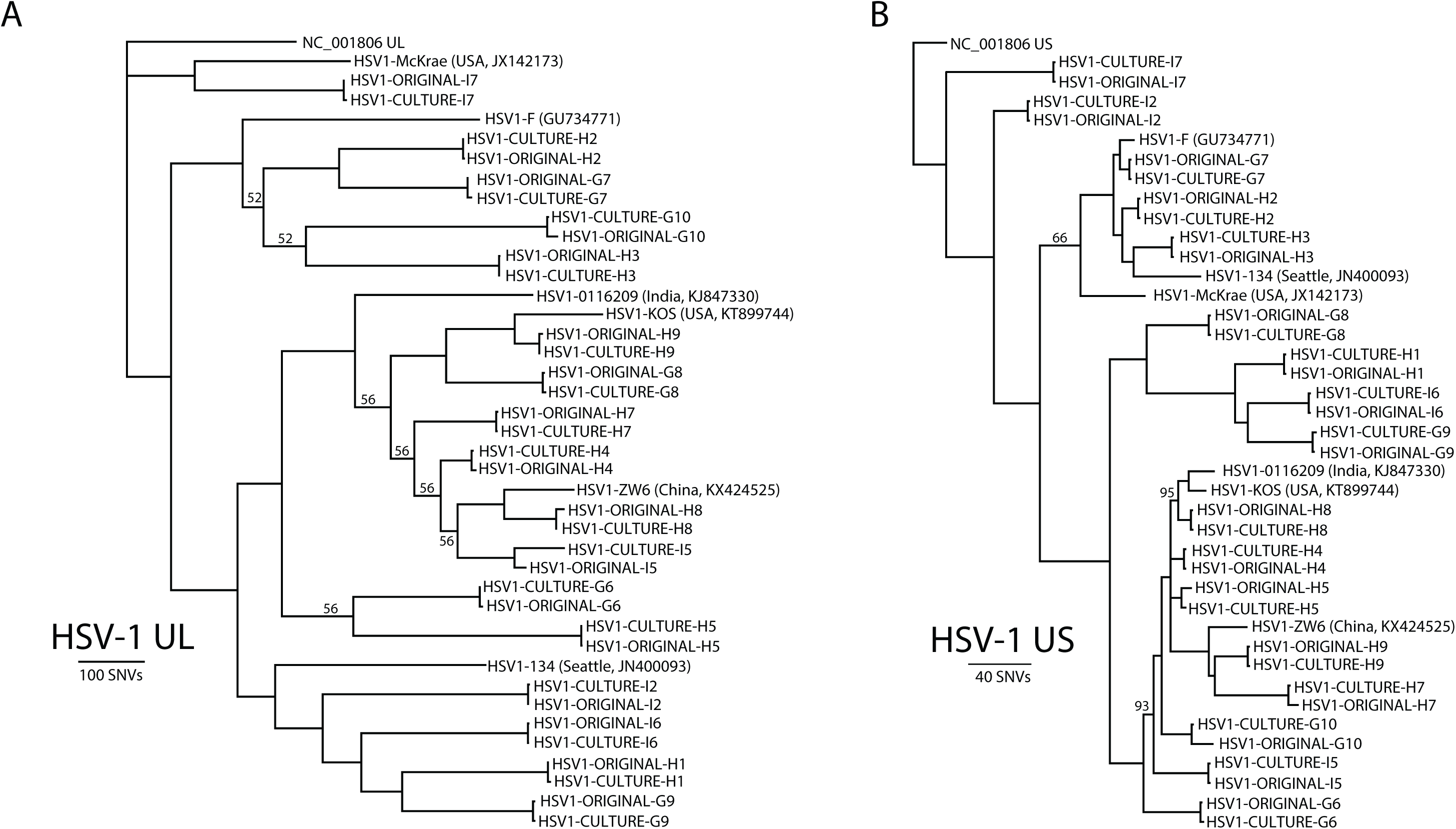
Limited evolution of HSV-1 during isolation in culture compared to sequence obtained directly from clinical samples. Phylogenetic analysis of UL (A) and US (B) sequences from HSV-1 subjected to capture sequencing after isolation in culture or directly from clinical sample. Across 14 of the paired samples, no single nucleotide variant was found in the UL or US region that was not present in homopolymers or UL36, US5, or US12 repeat regions. Of note, samples H4 and I5 were from the same patient 18 days apart illustrating HSV-1 oral superinfection. The long tree branch on I5 consensus sequence is due to changes in allele frequencies due to competitive viral growth in vitro between the superinfecting strains. All branch poster probabiltiies are >99% unless otherwise noted.

Sample G10 had four mutations between culture and clinical sample, including three synonymous changes in UL6, UL37, and UL54 and a T207A non-synonymous mutation in US7 coding sequence. All four mutations in G10 and the single mutation in H5 were confirmed by Sanger sequencing of the paired culture and original samples. The original sample for G10 had a notably low level of HSV-1 (18 copies/uL DNA or 9,000 copies/mL) and its assembly was 9.1% missing data (N’s). There was no evidence of HSV-2 recombination in the 17 pairs of HSV-1 sequences.

### Detection of HSV-1 superinfection

Samples pairs H4 and I5 were collected from the same patient in his 50s who underwent two allogeneic hematopoietic cell transplants for acute myelogenous leukemia. The first sample (H4, “day 1”) was collected from a tongue ulcer and the second sample (I5, “day 18”) was taken from an oral swab of a new tongue lesion 18 days later. He started foscarnet induction therapy 4 days prior to the first sample for treatment of CMV reactivation, but was not treated with acyclovir in the intervening period. The day 1 oral swab measured 10^5.9^ copies/mL for HSV-1 while the day 18 oral swab measured 10^5.4^ copies/mL.

After removing SNVs associated with the UL36 gene, the consensus UL sequences recovered from the two original oral swabs differed by 207 nucleotides, which is consistent with previous estimates of average pairwise SNV differences between two unrelated HSV strains (6, 9). The consensus UL sequence from the day 1 original sample and culture specimen differed at only 3 nucleotides, which were all associated with homopolymers, consistent with the lack of evolution seen during culture isolation for thirteen other paired HSV-1 specimens. However, the consensus UL sequence from the day 18 original sample and culture differed by 91 nucleotides, illustrating a rate of change significantly higher than seen in other paired specimens.

We hypothesized that changes in variant frequency between two different viral populations present in the day 18 specimen accounted for the increased rate of change during isolation in culture. Mapping of the day 18 original sample and cultured virus reads to the consensus day 1 original sample complete genome revealed 609 and 620 single nucleotide variants with minor allele frequency > 5% and depth > 25X. Most (92%) of the matched variant alleles increased in frequency from the original swab to the culture genome, from a median 45% to 66% allele frequency between the two specimens (Figure 5). These data suggest the difference in consensus genome between the culture and original day 18 specimens were due to allele frequency changes across the 50% consensus threshold within a mixed infection.

**Figure 5.**
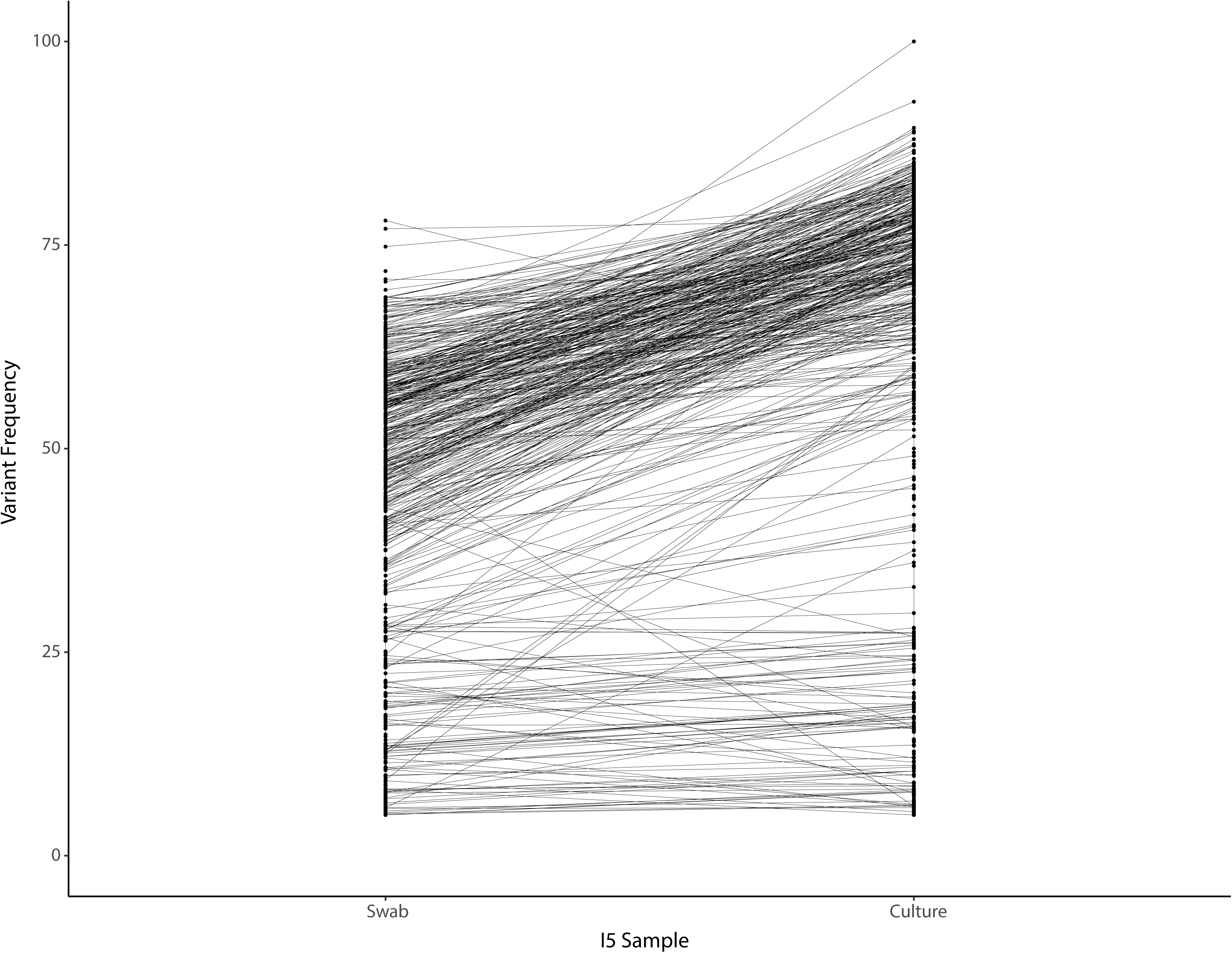
Allele frequency changes for the I5/”day 18” original oral swab HSV-1 genome and associated culture HSV-1 genome. The original consensus genome for the day 1 swab was used as a common reference from which to calculate allele frequency changes. The majority of alleles increase in frequency, crossing the 50% frequency threshold, resulting in artefactual evolution in culture that is the result of competition between mixed strains in culture.

Since the patient’s HSV-1 emerged during foscarnet therapy, we next interrogated our sequence data for whether antiviral resistance was present in either of the oral swabs. Four non-synonymous mutations were present in the UL30 gene from the day 18 oral swab compared to the day 1 oral swab at varying allele frequencies (S724N, 6%; E798K, 11%; I810L, 16%; F918L, 57%). Compared to the HSV-1 17 strain reference genome (NC_001806), both original samples had consensus UL30 coding changes at S33G, V905M, P920S, P1199Q, and T1208A. None of these changes has previously been reported to be associated with foscarnet resistance (27–29). These results are consistent with the patient being superinfected with two separate HSV-1 strains that reactivated at separate times on the patient’s tongue and were simultaneously detected from the day 18 specimen. These data also indicate that in the setting of superinfection, cultured samples may appear very different from swab samples due to differential abilities of the multiple viruses to grow in culture.

## Discussion

We report the validation of capture sequencing panels for obtaining near complete HSV-1 and HSV-2 genomes directly from clinical samples. The panels allow the recovery of HSV-1 and HSV-2 genomes in approximately 3-5 days with as few as 100,000 paired-end reads at viral concentrations that are up to 100,000-fold lower than previously reported for herpes simplex viruses. The level of enrichment seen here is similar to seen by others using capture panels (30, 31). We used this panel to show that HSV-1 undergoes extraordinarily limited evolution during culture isolation, finding only 5 single nucleotide variant across more than 1.8 megabases of UL-US sequence from 15 paired HSV-1 positive samples.

To date, direct-from-sample whole genome sequencing for herpes simplex viruses has been limited to samples with extraordinarily high viral copy numbers (9). The vast majority of genome sequence data available from herpes simplex viruses comes from culture isolates. Our data indicate that these culture isolate sequences likely faithfully represent the original herpes simplex virus sequence present in the clinical samples from which the viral isolate originated.

Despite the success of culture in faithfully amplifying genomes, capture sequencing direct from patient samples has a number of advantages. Clinical samples at low HSV copy number often do not yield positive cultures. Even high copy number culture samples may consist of less than 1% of HSV-1 reads, and thus captured libraries can be sequenced in greater depth and in a more multiplexed fashion. Capture sequencing of HSV for genotypic antiviral resistance for drugs such as acyclovir or foscarnet may also return results back faster than phenotypic culture-based tests, which require growth of the virus and have a relatively long turn-around time. Though this assay can be performed in as little as 72 hours, we envision that a capture-based whole-genome genotypic clinical test for antiviral resistance or epidemiological purposes would likely be batched weekly with a sample-to-answer turn-around time of 5-11 days, pending when the sample is received and the required test volume. Engineering and automation improvements to the protocol could substantially reduce hands-on-time and lead to significantly faster turn-around times.

We also use direct from sample sequencing to show the first case of HSV-1 superinfection detected directly from a patient by next-generation sequencing. The prevalence of HSV-1 infected individuals that carry more than one HSV-1 strain is not known, while HSV-2 superinfection is estimated to occur in approximately 3.5% of patients positive for HSV-2 (10). Despite HSV-1 reactivating in this patient in the setting of foscarnet treatment, no previously characterized mutations for foscarnet resistance were discovered (27, 29, 32). These data underscore the current challenge in confidently assigning antiviral resistance for HSV through genomic sequence.

Limitations of our study include examining HSV-1 evolution in the context of brief culture exposure with minimal passage. Our results may not be reflective for strains that undergo more passages than the initial viral isolation (“zero passage”) that was performed here. Notably, we and others have also not solved the problem of the high degree of homopolymers and repetitive sequence in the setting of high GC content in human herpes simplex viruses. Indeed, several of the loci cannot be confidently synthesized as oligonucleotides for the affinity purification panel. We also limited our sequence analysis to the UL and US regions of the genome.

In summary, we demonstrate the validation of a new robust, accurate, and sensitive tool to recover near complete HSV-1 and HSV-2 genome sequences, along with an easy pooling scheme to reduce overall sequencing costs. We show that HSV-1 culture isolates undergo very few genomic changes in the UL-US region during isolation in culture. Indeed, culture may be the ultimate viral enrichment method for HSV-1 and HSV-2.

## Materials and Methods

### Clinical Samples

HSV-1 and HSV-2 samples were selected from natural history research studies at the University of Washington Virology Clinic that spanned a range of pre-capture viral concentrations (Table S1). Excess HSV-1 samples sent to the University of Washington Clinical Virology Lab for culture over a one-month period in 2017 were also selected for sequencing (Table S2). Informed consent was obtained for HSV-1 and HSV-2 specimens from the Virology Clinic. Informed consent was waived for HSV-1 original swab and culture evolution samples by the University of Washington Human Subjects Division based on use of de-identified excess HSV-1 clinical specimens. The University of Washington Human Subjects Division approved both procedures.

### Swab DNA extraction and qPCR

DNA was extracted from 200 µl of proteinase K buffer that the original swab specimen was placed in or from 40 µl of viral culture supernatants using QIAamp DNA Blood Mini kit (Qiagen). DNA was eluted into 100 ul of AE buffer provided in the extraction kit and 10ul of the DNA was then used for each real-time PCR reaction. HSV DNA copy number was measured by a HSV type common real-time PCR assay which amplifies the gB gene (33). Human genomic number in the original swab samples was measure by the primers and probe design to detect beta-globin gene (betaF: TGA AGG CTC ATG GCA AGA AA; probe: TCC AGG TGA GCC AGG CCA TCA CT; betaR: GCT CAC TCA GTG TGG CAA AGG). Each 30 ul PCR reaction contained 10 ul of purified DNA, 833 nM primers, 100 nM probe, internal control, 15 ul of QuantiTec multiplex 2x PCR master mix. The thermocycling conditions were as following: 50°C for 2 minutes, 95°C for 15 minutes, and followed by 45 cycles of 94°C for 1 minute and 60°C for 1 minute.

### PCR/Sanger sequencing

PCR reactions of HSV-1 and HSV-2 UL23 genes and discrepant loci were performed using the PrimeSTAR GXL DNA Polymerase (Takara) with the primer sequences available in Table S3A/B. Each 50 ul PCR reaction contained: 10 ul DNA, 10 ul 5X PrimerSTAR GXL buffer, 0.2 mM dNTP, 0.32 µM primers, and 1.25 units of PrimeSTAR GXL DNA Polymerase. PCR reactions were performed using the following conditions (98C 45s, [98C 10s, 60C 15s, 68C 120s] x 40 cycles, and 68C 10min). Confirmatory PCR for discrepant loci was performed using the following conditions (98C 415s, [98C 10s, 55C 15s, 68C 30s] x 40 cycles, and 68C 5min). Sanger sequencing reactions were performed using the sequencing primers in Table S3C.

### Capture sequencing of HSV-1 and HSV-2 samples

We first optimized the fragmentation and library preparation steps on high concentration HSV-2 culture specimens, comparing Nextera XT, Kapa HyperPlus, and custom NEB Fragmentase-based protocols. Kapa HyperPlus and NEB Fragmentase gave equivalent coefficients of variation for genome coverage (29.2% versus 31.0%), while Nextera XT coefficient of variation was three times higher (96.7%), likely due to the known GC bias of the enzyme (data not shown). We subsequently chose to perform pre-capture library preparation using half-volumes of Kapa HyperPlus with a 7-minute fragmentation step on 100ng of DNA, ligation of 15uM common Y-stub adapters, and 0.8X Ampure post-ligation cleanup. Post-loigation PCR amplification was performed using the Kapa HiFi HotStart ReadyMix with Truseq dual-indexed primers (98C 45s, [98C 15s, 58C 30s, 72C 30s] x 12 cycles, and 72C 1 min) and cleaned using 0.8X Ampure beads. Pre-capture libraries are quantitated on a Qubit 3.0 Fluorometer (ThermoFisher).

Prior to capture, libraries were pooled in sets of 4-10 libraries based on the ratio of HSV-1/2 to beta-globin and total number of HSV-1/2 copies present in each library. A total of 300-500ng DNA is targeted for each pool. We aim for less than 10-fold variance from the highest to lowest concentration in HSV-1/2 copies within each pool. Hybridization capture is performed according to the IDT xGen protocol (version 2). Capture panels were designed as 1X tiling 120-bp panels according to HSV-1 strain 17 and HSV-2 HG52 reference sequences (NC_001806, NC_001798) with human masking based on IDT xGen design. Oligonucleotide capture panel sequences are available in DataSet S1/S2.

### Computational pipeline for assembly and annotation of HSV genomes

Our workflow combines multiple open source tools with custom shell and R scripts to rapidly extract and annotate near-full length HSV genomes from raw Illumina sequencing reads (Figure 2). All code is available on Github (https://github.com/proychou/HSV).

Raw sequencing reads (either paired or single-end) in fastq format are trimmed to remove adapters and low-quality regions using BBDuk (34). QC reports are generated on the raw and preprocessed files using FastQC (35). Optionally, non-HSV reads are filtered out using BBduk with k = 31 and hdist = 2. Preprocessed reads are *de novo* assembled using SPAdes and contigs are ordered by aligning to HSV-1 or −2 reference sequences (NC_001806, NC_001798, KF781518) using Mugsy (36, 37). A custom script in R/Bioconductor is used to fill in any gaps between contigs to create a template and reads are mapped to this template using Bowtie2 (38). A second script using R/Bioconductor is used to construct and clean up the final consensus sequence and prepare files for annotation. Annotation is performed using Prokka and a custom script to construct the final consensus sequence (39).

Although designed to be run on a high-performance computing (HPC) cluster, the code can also be run on a desktop computer. Additional wrapper scripts are available for parallelization of samples on an HPC cluster with scheduling systems like SLURM or PBS/Torque. Consensus sequences for each pair were aligned using MAFFT, pairwise differences calculated, UL-US extracted and locations of differences determined by adding annotations from HSV 1 references. The ggplot2 and nplr packages were used in R to calculate the limits of genome recovery (40). Phylogeny were created using MrBayes with default parameters (41).

### Recombination analysis of HSV-1 culture

HSV-1 isolate sequences were examined individually for HSV-2 recombination using alignment trios with Chimp HSV (NC_023677.1) and an HSV-2 reference sequence (KF781518.1) as input for RDP (version Beta 4.95). The RDP program was run from the command line with the default settings (42). This program uses the RDP, GENECONV, Chimaera, and MaxChi algorithms to both detect events and verify events identified by other algorithms. The algorithms BootScan, SiScan, and 3Seq are computationally intensive when used to detect new events and so are only used to verify other events when using the default settings. All output files were combined and screened for p-value < 1×10^−10^ for at least three algorithms. Results were the same when all putative events having a p-value of 1×10^−10^ or smaller for only 2 algorithms were considered.

### Culture of HSV-1 isolates

Swab samples were collected and transported to the clinical lab in universal transport medium. Supernatant fluid was removed, diluted with Hanks Balanced Salt Solution (HBSS) with antibiotics, centrifuged at 700xg for 10 minutes, and 0.2 mL was inoculated into duplicate Human Fibroblasts (MRHF) (Diagnostic Hybrids). Cell monolayers were observed microscopically daily for HSV cytopathic effect (CPE). If typical CPE noted, culture media was harvested and frozen at −80°C for PCR analysis. To confirm subtype of isolate, MRHF cells were scrapped and spotted onto welled slides, air-dried, fixed in acetone and stained with monoclonal antibody to HSV-1 and HSV-2 (MicroTrak, Trinity Biotech).

## Acknowledgements

AW has received funding for clinical trials from Genocea and Vical through the University and consulting fees from Aicuris.

## Supplemental material

**Figure S1** – HSV-1 culture isolate captured with HSV-2 capture panel. Early in development, we attempted capture of an HSV-1 culture isolate with an HSV-2 capture panel. A) Coverage map of reads across the HSV-1 genome shows coverage was poor. Despite an average coverage of 179X, only 58% of the HSV-1 UL region had depth >= 10X. Y-axis denotes read depth, while X-axis is the genome position for HSV-1. HSV-1 genes are denoted in green while repeat regions are highlighted in light blue. B) HSV-1 UL locus depth correlates with pairwise identity to HSV-2 UL sequence. We calculated the pairwise HSV-1 versus HSV-2 sequence identity across a 120 nucleotide sliding window and plotted as a histogram (blue). For each 120 nucleotide bin across the HSV-1 UL we calculated the median bin depth from the capture sequencing normalized to the maximum bin depth (black dots).

**Figure S2** – HSV-1 (A) and HSV-2 UL23 (B) genes show 100% identity whether sequenced by PCR-Sanger or capture panel next-generation sequencing approach.

Table S1A. HSV-1 capture efficiency

Table S1B. HSV-2 capture efficiency

Table S2A. HSV-1 swab and culture sequencing metadata

Table S2B - History HSV1+ clinical samples in culture. Cytopathic effect was checked every day and graded on a scale of 0-4 before harvest. H=Harvest

Table S3A – PCR primers for U23

Table S3B – Primers for PCR and confirmatory Sanger sequencing of discrepant original swab versus culture samples

Table S3C – Sequencing primers for U23

DataSet S1. Capture panel design for HSV-1

DataSet S2. Capture panel design for HSV-2

